# FISHNET: A Network-based Tool for Analyzing Gene-level P-values to Identify Significant Genes Missed by Standard Methods

**DOI:** 10.1101/2025.01.29.635546

**Authors:** Sandeep Acharya, Vaha Akbary Moghaddam, Wooseok J. Jung, Yu S. Kang, Shu Liao, Michael A. Province, Michael R. Brent

## Abstract

FISHNET uses prior biological knowledge, represented as gene interaction networks and gene function annotations, to identify genes that do not meet the genome-wide significance threshold but replicate nonetheless. Its input is gene-level P-values from any source, including omicsWAS, aggregation of GWAS P-values, CRISPR screens, or differential expression analysis. It is based on the idea that genes whose P-values are low due to sampling error are distributed randomly across networks and functions, so genes with suggestive P-values that cluster in densely connected subnetworks and share common functions are less likely to reflect sampling error and more likely to replicate. FISHNET combines network and function analysis with permutation-based P-value thresholds to identify a small set of exceptional genes that we call FISHNET genes.

Applied to 11 cardiovascular risk traits, FISHNET identified 19 gene-trait relationships that missed genome-wide significance thresholds but, nonetheless, replicated in an independent cohort. The replication rate of FISHNET genes matched or exceeded that of other genes with similar P-values. FISHNET identified a novel association between *RUNX1* expression and HDL that is supported by experimental evidence that *RUNX1* promotes white fat browning, which increases HDL cholesterol levels. FISHNET also identified an association between *LTB* expression and BMI that is supported by experimental evidence that higher LTB expression increases BMI via activation of the LTβR pathway. Both associations failed genome-wide significance thresholds, highlighting FISHNET’s ability to uncover meaningful relationships missed by traditional methods. FISHNET software is freely available at https://doi.org/10.5281/zenodo.14765850.

## Introduction

The primary goal of hypothesis testing is to determine whether an observation made in a sample from a population is likely to be true in the population. Statistical hypothesis testing is generally used to minimize the risk of false positive findings at the cost of substantial risk of false negatives. The false-negative risk is magnified by multiple testing correction when many tests are carried out, as is the case in genome-wide association studies (GWAS) [1, 2], transcriptome-wide association study (TWAS) [3, 4], and other omics-WAS [5, 6]. Common methods of correcting for multiple testing include Bonferroni adjustment [7, 8] and Benjamini-Hochberg (BH) false discovery rate (FDR) control [9, 10]. Moreover, when effect sizes are small, as is typical in genetics, very large samples are needed to reach statistical significance. Thus, there is an urgent need for methods that can identify true, population-wide omics-trait relationships that do not meet Bonferroni or BH significance thresholds. Such methods can be evaluated empirically by replication studies, in which additional samples are drawn from the population.

We propose a new method for finding exceptional gene-trait associations that replicate at a higher rate than other genes with the same adjusted p-values, which we call FISHNET (Finding Significant Hits in Networks). FISHNET integrates results from gene-level summary statistics with prior biological knowledge represented as networks. It can use gene-level summary statistics from GWAS, TWAS with measured or predicted gene expression levels, proteome-wide association studies, RNA-Seq experiments, functional genetics screens, or any other source. It uses gene-gene interactions from co-expression networks, protein-protein interaction (PPI) networks, or other networks, together with gene function annotations from Gene Ontology (GO) [11]. We hypothesized that genes whose p-values are low due to sampling error are distributed randomly across biological networks and functions. Thus, when genes with low p-values cluster in densely interconnected subnetworks (network modules) and share common functions, they are less likely to reflect sampling error and therefore more likely to replicate in new samples. FISHNET combines network module enrichment analysis [12] and GO over-representation analysis [13] with permutation-based significance thresholds to identify a small set of exceptional, trait-influencing genes that we call FISHNET genes.

FISHNET is specifically designed to identify gene-trait associations that replicate in independent samples at a higher rate than other associations with similar p-values. In addition to prioritizing genes that meet traditional significance thresholds, it identifies genes below these thresholds that replicate at a similar rate, enabling thresholds to be relaxed. Notable features include its permutation-based significance testing, its automated testing of model assumptions, and its use of GO biological process annotations in addition to networks. These annotations, which are manually curated and reflect a large body of literature [11], are complementary to networks, which are typically generated from high-throughput data. Another reason for supplementing networks with GO analysis is that network modules are identified computationally [14], based on network connectivity, so there is no guarantee that two genes in the same module share a common function.

This paper makes four contributions. First, it introduces the FISHNET algorithm and software. The software is freely available as a Docker container, making it easy to install and deploy across a wide range of computing environments. Second, it evaluates the replicability and reproducibility of FISHNET genes using 43 sets of GWAS gene-level summary statistics [14]. Evaluation was carried out on nine combinations of networks and algorithms for finding modules, identifying the most useful combinations. Third, using the best networks and the best module detection algorithm, it identifies FISHNET genes across 11 traits associated with cardiovascular risks in the Long Life Family Study (LLFS) cohort and performs the replication analysis in the Framingham Heart Study (FHS) cohort [15, 16]. The results show that FISHNET genes have a better replication rate than non-FISHNET genes with similar p-values. Fourth, this paper introduces metrics to assess whether user inputs – network modules and gene-level summary statistics – violate the model assumptions.

The software can be accessed at https://doi.org/10.5281/zenodo.14765850.

## Methods

### Gene-level Summary of GWAS

Gene-level summary statistics were obtained from 185 meta-analyses of GWAS collected for the 2019 Disease Module Identification DREAM Challenge [14]. The SNP-level summary statistics were aggregated to gene-level statistics using PASCAL [12]. PASCAL uses the sum of chi-squared approach to calculate a gene-level p-value. To create discovery and replication set pairs for replication analysis, we used only traits that had more than one study with completely or partially independent cohorts. Additionally, we removed studies where the genotyped SNPs did not cover all chromosomes in the genome. After these exclusions, we retained gene-level summary statistics from 43 GWAS. There were 17 traits with exactly two GWAS datasets – Bipolar disorder, BMI, BMI (male), BMI (female), Coronary artery disease, Fasting glucose, Height, Hip circumference (male), Hip circumference (female), Molecular degeneration, Rheumatoid arthritis, Total cholesterol, Waist circumference (male), Waist circumference (female), Waist-hip ratio (male), Waist-hip ratio (female), and Schizophrenia. For High-density lipoprotein (HDL), Low-density lipoprotein (LDL), and Triglycerides, there were three GWAS studies, two of which we used as discovery sets, with the third as the replication set for both. The final dataset consisted of 23 discovery and replication set pairs, each with gene-level GWAS summary statistics. The characteristics of each dataset can be found in **Supplementary Table 1**.

### Long Life Family Study (LLFS)

The Long Life Family Study (LLFS) is a longitudinal family study that enrolled families enriched for exceptional longevity to discover genetic factors contributing to healthy aging. LLFS enrolled 4,953 participants in 539 pedigrees, primarily of European ancestry (99%). The recruitment procedure and enrollment criteria of the LLFS participants have been previously described [17, 18]. The data generated in the study includes gene expression levels from blood and biomarkers of health and aging. We focused on 11 traits associated with cardiovascular risks spanning four categories: pulmonary (forced expiratory volume, forced vital capacity, and the ratio of the two), lipids (high-density lipoprotein, low-density lipoprotein, triglycerides, total cholesterol), anthropometric (BMI, BMI-adjusted waist), and cardiovascular (pulse, ankle-brachial index) [19-23].

### LLFS RNA-Seq and Transcriptome Wide Association (TWAS)

RNA extraction and sequencing were carried out by the McDonnell Genome Institute at Washington University (MGI). Total RNA was extracted from PAXgene(tm) Blood RNA tubes using the Qiagen PreAnalytiX PAXgene Blood miRNA Kit (Qiagen, Valencia, CA). The RNASeq data were processed with the nf-core/RNASeq pipeline version 3.3 using STAR/RSEM and otherwise default settings (https://zenodo.org/records/5146005). The RNA-Seq data QC steps and the gene expression level adjustment model used in this study have been previously described by Acharya et al., [4] who performed TWAS on the same 11 traits using the data from the first clinical exam in LLFS. Since our previous publication, the dataset has grown. Depending on the trait, the number of participants with both RNA-Seq and trait data now ranges from 879 to 1667. The adjustment steps for all 11 traits are as described [4]. **Supplementary Table 2** shows the characteristics of study participants for covariates and 11 cardiovascular traits.

For each trait, the adjusted gene expression residuals were used as a predictor and the adjusted trait residuals were used as a response variable in a linear mixed model implemented in MMAP [24]. A kinship matrix generated by MMAP from the LLFS pedigree was used to account for family relatedness. For traits with genomic inflation factor (GIF) > 1.1, the p-values were adjusted by using BACON [25]. The same RNA-Seq processing and trait adjustment steps were applied for replication in the Framingham Heart Study (FHS) dataset, where the number of participants with data for each trait ranged from *n =* 1080 to *n =* 1380.

### Module enrichment analysis and Gene Ontology (GO) Over-representation Analysis

First, in each selected gene-gene interaction network, network modules (highly connected subnetworks) were identified by three algorithms from the Disease Module Identification DREAM Challenge [14], designated as R2, based on random walk, K1, based on kernel clustering, and M2, based on modularity optimization. The selected modules were from STRING functional protein-protein Interaction (PPI) network [26], InWeb physical PPI network [27], and a gene co-expression network from Gene Expression Omnibus [28, 29]. After identifying network modules, GWAS or TWAS gene-level summary statistics and sets of genes in network modules were fed into PASCAL’s module enrichment algorithm [12]. Figure 1A depicts the inputs and outputs of module enrichment analysis and GO Over-representation analysis. After modules were identified, the connections between genes in modules were not used. PASCAL’s module enrichment algorithm outputted sets of genes in modules significantly enriched for genes with low p-values, adjusted for the total number of modules tested using Bonferroni correction. GO over-representation analysis was done on the set of genes in each enriched module by using WebGestaltR package (version: 0.4.6) with the following configuration: (organism: hsapiens, method: ORA, enrichDatabase: GO Biological Process, FDRMethod: BH, FDRThreshold = 0.05) [13]. The affinity propagation feature in WebGestaltR was used to eliminate GO biological processes with highly overlapping member genes, thereby reducing the multiple testing burden and improving computational performance.

**Fig 1:**
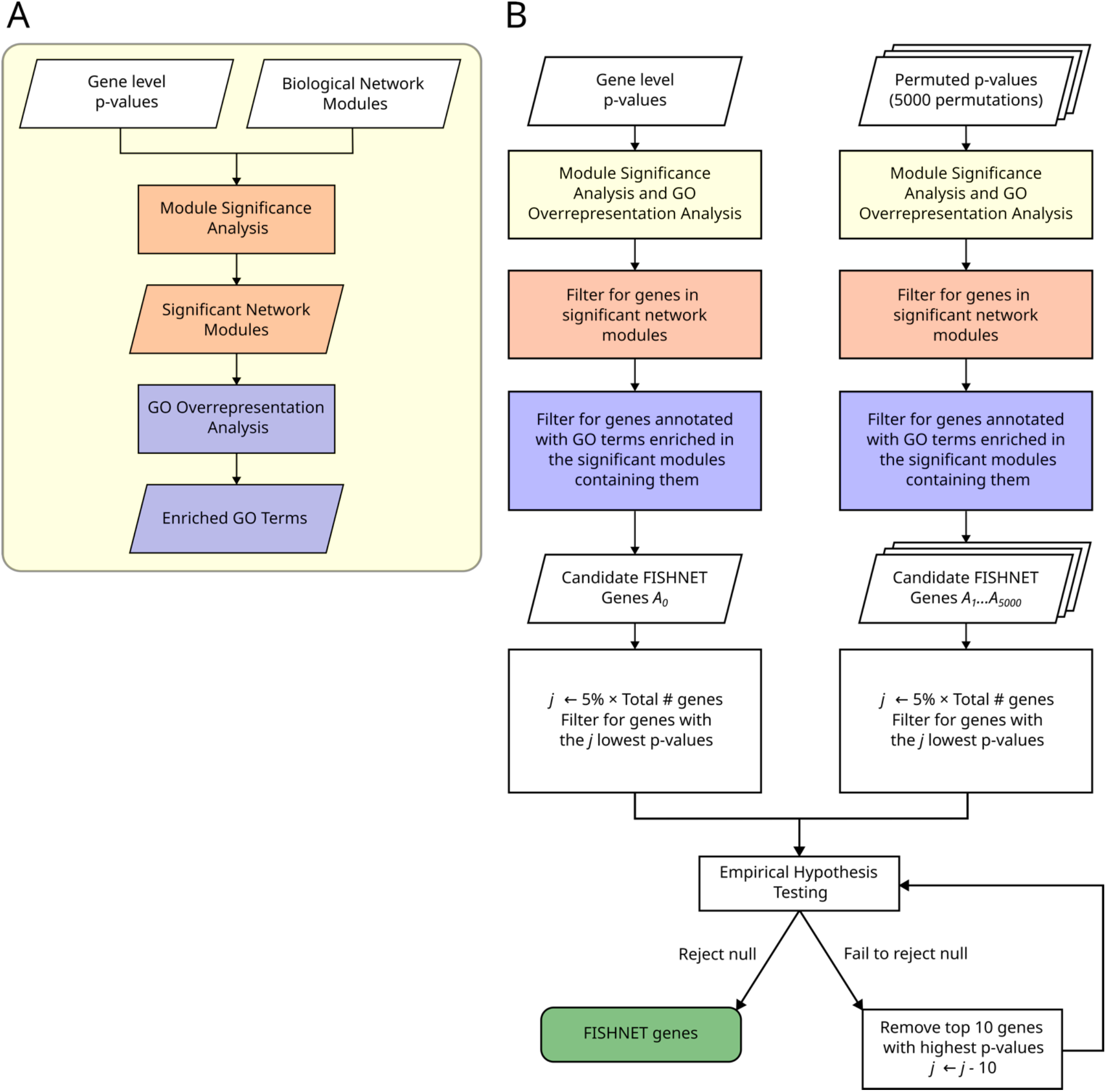
(A) The gene-level p-values are input into module significance analysis. Module significance analysis outputs significant modules and their p-values. Gene ontology over-representation analysis identifies biological processes with significant over-representation among genes in each significant module. (B) The workflow illustrates the gene prioritization mechanism used to identify FISHNET genes. For details of the permutation-based hypothesis testing, see Methods.

### FISHNET Algorithm

Fig 1b depicts the FISHNET algorithm. FISHNET is run separately for each combination of traits, networks, and module detection algorithms. Genes are considered caught in the FISHNET if they:

1. Are in a network module unusually enriched for genes with low p-values (PASCAL’s module enrichment analysis). This passes only genes that work together with other trait-implicated genes.
2. Are annotated by a gene ontology (GO) biological process term that is enriched in their module (GO Over-representation analysis). This passes only genes that work together as a part of a common biological process.
3. Are among the top *N* genes ranked by p-values. The permutation-based null model described below determines *N. N* is constrained to be, at most, 5% of genes. If all p-values are under the null model, they should uniformly distributed, so the top 5% corresponds to a nominal p-value threshold of 0.05; if the p-value distribution is inflated, which is common in TWAS with measured gene expression levels, the top 5% will correspond to a p-value threshold lower than 0.05.

### Permutation-based null model: Empirical Hypothesis Testing

Null hypothesis: Among candidate FISHNET genes, we test the null hypothesis that the number of genes that (1) lie in significant modules and (2) participate in biological processes enriched among module genes is not substantially higher than expected at random.

PASCAL’s module enrichment analysis pipeline monotonically transforms the genes’ p-value distribution into a chi-squared distribution. This process relies only on the ranks (quantiles) of the genes when ranked by their p-values. Therefore, FISHNET works as follows:

1. Randomly permute the original p-value ranks of genes 5000 times.
2. Run module enrichment and GO over-representation analyses for the original gene ranks and all permutations.
3. Let *A*_*i*_ be the set of genes in permutation *i* that:
  a. Are in a module of interacting genes that is significantly enriched for low p-values.
  b. Are annotated with a biological process term enriched among genes in that module. The set of genes satisfying these criteria obtained from the true ranks is referred to as *A*_0_ – candidate FISHNET genes.
4. Let *K* be 0.05 times number of genes analyzed. Rank genes by gene-level p-value, from smallest to largest, and let B_*j*_ be the set of genes within p-value rank *j*, for *j* = 10 to K (inclusive) by 10s. For example, B_*30*_ is the set of 30 genes with the smallest gene-level p-values. Let C_*ij*_ = A_*i*_ *∩* B_*j*_, where *i* ranges from 1 to 5000 for random permutations and *i =* 0 for original ranks.
5. The expected FDR for genes above rank j, FDR, is 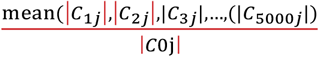
6. Let PercentileRank_*j*_ be the quantile of |C_0*j*_| in |C_1*j*_|, |C_2*j*_|, |C_3*j*_|, …., |C_5000*j*_|, with larger quantiles corresponding to larger sets.
7. *N =* max(*j*) such that FDR_*j*_ <= 0.05 and PercentileRank_*j*_ >= 99%.
8. Set of FISHNET genes = A_*0*_ ∩ B_*N*_ = C_*0N*_.

## Results

We developed the FISHNET algorithm to identify replicable gene-trait relationships missed by standard association analyses. It works by combining gene-level summary statistics from association analyses with prior biological knowledge encoded in gene-gene interaction networks and gene functional annotations. Among the genes with suggestive/significant p-values from association tests, FISHNET prioritizes genes that cluster in network modules with other genes that have the same functional annotations similarity (See Fig. 1 and Methods). We applied FISHNET to GWAS gene-level summary statistics from Choobdar et al., 2019 [14] (**Supplementary File 1, Supplementary Table 1**) and TWAS summary statistics from the LLFS and FHS cohorts (**Supplementary File 2**) to test whether FISHNET recovers replicable gene-trait associations missed by traditional significance thresholds.

### FISHNET performance varies across networks and modularization algorithms

FISHNET was applied to 23 GWAS discovery-replication summary statistics pairs and gene sets from network modules in the STRING functional PPI network [26], InWeb physical PPI network [27], and gene co-expression network [28, 29]. Three algorithms were used to identify modules in each network. The algorithms are based on three different approaches: kernel clustering, modularity optimization, and random walk. Thus, we ran FISHNET 9 times on each discovery-set replication set pair. Each FISHNET output gene (hit) from each run can be uniquely identified by four factors: the gene, network, modularization algorithm, and trait used in the run. For convenience, we refer to these unique identifiers as *genequads* (Fig 2a). Networks were evaluated based on the reproducibility and replicability of all gene quads based on that network (unioned over the other three factors). Likewise, modularization algorithms were evaluated on all genequads based on that algorithm, unioned over other factors. A FISHNET quad from a discovery set is reproduced if it is also a FISHNET quad in the replication set. A FISHNET quad is replicated if a hit identified in the discovery set is Bonferroni significant in the replication set.

**Figure 2:**
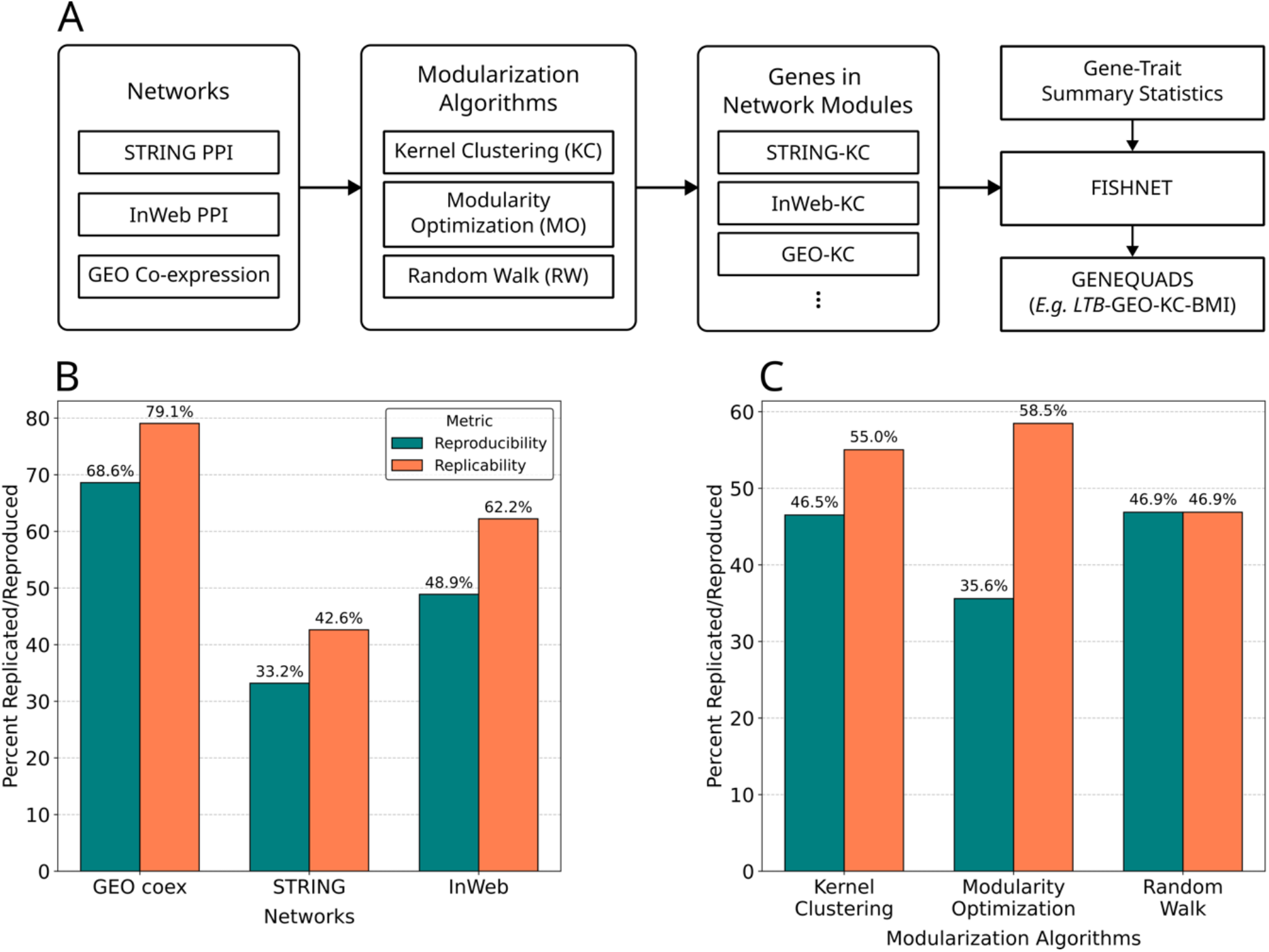
Replication and reproducibility rates across networks and modularization algorithms using 23 pairs of GWAS discovery and replication summary statistics. (A) Three modularization algorithms are applied to three networks to obtain nine sets of genes, each fed into FISHNET with gene-trait summary statistics. (B) GEO co-expression outperforms other networks in replicability and reproducibility, while STRING functional PPI performs worst in both metrics. (C) Kernel Clustering achieves the best balance of replication and reproducibility, while Modularity Optimization performs best in replication, and Random Walk performs best in reproducibility.

Across 23 GWAS discovery datasets, three networks, and three modularization algorithms, FISHNET identified 375 GeneQuads, of which 162 (43.2%) reproduced and 200 (53.3%) replicated in replication sets (Supplementary Table 3). Among networks, the GEO co-expression network had the best reproducibility and replication rate, followed by InWeb PPI (Fig 2a). Among modularization algorithms, random walk had the best reproducibility rate and modularity optimization had the best replication rate. Kernel clustering-based modularization algorithm had the best performance when both replicability and reproducibility were considered. The rest of the analyses discussed in this paper are based on the two best-performing networks, InWeb PPI and GEO co-expression, and the single best modularization algorithm, kernel clustering.

### FISHNET identifies replicable gene-trait relationships missed by association analyses

We previously published results from association analyses of blood gene expression levels with 11 cardiovascular risk traits in the LLFS cohort. For the current paper, we applied FISHNET to those summary statistics to identify replicable gene-trait relationships that did not reach significance thresholds. Across 11 traits, FISHNET identified 287 unique gene-trait relationships, 34 of which replicated in the FHS cohort. Nineteen of the 34 were not Bonferroni significant in the LLFS cohort (Table 1). For pulse and Forced Vital Capacity (FVC) there were no significant hits in the original study, but both have one replicated FISHNET gene. For BMI, High-Density Lipoprotein (HDL), and Triglycerides, FISHNET identified 15 replicated gene-trait relationships that were genome-wide significant in the original studies and 17 that were not.

**Table 1:** FISHNET identified replicated gene-trait relationships across five traits in LLFS, including both genes that meet the genome-wide significance threshold and those that do not.

| Genes | Trait | Bon. Significant in LLFS ($p \leq 3.5 \times 10^{-6}$) |
| --- | --- | --- |
| *PTGER2, FFAR2, CX3CR1, HCK, BIRC3, UBE2J1, DUSP2* | BMI | Yes |
| *TNF, LTB, FLT3, CD22, CLEC4E, MMP8, TLR5, CCR6, SERPINA1, EDA, IL18R1, PEA15* | BMI | No |
| *MS4A3, SREBF2, GATA2* | HDL | Yes |
| *CD180, GCA, RUNX1* | HDL | No |
|  | Pulse | Yes |
| *ITGAM* | Pulse | No |
| *TCN1, MS4A3, SREBF2, RUNX1, GATA2* | Triglycerides | Yes |
| *LTB, CD244* | Triglycerides | No |
|  | FVC | Yes |
| *CD1D* | FVC | No |

FISHNET identified an association between expression of *LTB*, the gene encoding lymphotoxin-beta, and BMI, which is strongly supported by experimental studies in mice. *LTB*, a member of the tumor necrosis factor family, regulates immune responses through the activation of the LTβR pathway [30, 31]. Inactivation of this pathway in Ltβr^-/-^ mice confers resistance to diet-induced obesity, possibly through its effects on the gut microbiota [32]. The direction of this effect is consistent with our finding that expression of *LTB* is positively associated with higher BMI in humans. The mouse experiment supports the possibility that higher expression of *LTB* increases BMI by activating the LTβR pathway. Given that cytokine-mediated immune responses are key mechanisms in obesity-induced inflammation [33, 34], obesity may also induce expression of *LTB* in a positive feedback loop. One previous study showed upregulation of *LTB* in peripheral blood monocytes of 14 mildly obese Korean men but not in 12 moderately obese men [35]. However, there is no previous evidence for association between the expression level of *LTB* and BMI in the TWAS Atlas [36], nor is there any previous genetic evidence linking *LTB* to BMI in humans in the GWAS Catalog [37]. Although *LTB* did not reach genome-wide significance for BMI in our TWAS, FISHNET supported an association because: (1) *LTB* is in the same module of the co-expression network with other genes that show evidence of association with BMI, including *SERPINA1, HCK*, and *CX3CR1*, and (2) *LTB* is annotated with GO terms involving cytokine production that are over-represented among genes in that module. There is no guarantee that genes with similar expression patterns will be involved in the same molecular processes, but the combined FISHNET criteria highlighted a relationship between *LTB* and CX3CL1 that is supported by biological evidence -- LTB/LTβR signaling induces expression of the pro-inflammatory chemokine CX3CL1, a *CX3CR1* ligand [38].

FISHNET also identified a novel, positive association between *RUNX1*, the gene encoding the Runt-related transcription factor 1, and HDL. This finding is strongly supported by experimental evidence highlighting *RUNX1*’s role in promoting white fat browning, which in turn increases HDL cholesterol levels. CDK6 negatively regulates white fat browning by suppressing *RUNX1* expression, which directly promotes the expression of key thermogenic genes such as *Pgc-1α* and *Ucp-1* [39]. RUNX1 protein promotes white fat browning by binding to the promoters of the brown-adipose-tissue-specific genes *Pgc-1α* and *Ucp-1* in inguinal white adipose tissue (iWAT) [39]. In an experiment aimed to understand the role of *RUNX1* in mediating the effects of CDK6 protein on white-fat browning, deletion of *RUNX1* in mice lacking CDK6 reduces the expression level of *Pgc-1α* and *Ucp-1*, consequently inhibiting white-fat browning[39]. Furthermore, inducing white fat browning via Kaempferol (KPF) treatment in mice under both a high-fat diet and normal chow diet enhances RUNX1 protein levels and increases the expression of *Pgc-1α* and *Ucp-1* in iWAT *[40]*. Chemical induction of white fat browning via KPF treatment has also been directly linked to the change in HDL-cholesterol levels. Specifically, when white fat browning is induced in mice with hypertriglyceridemia (Apoa5-/-), the cholesterol shifts from the triglycerides-rich lipoproteins to HDL, increasing the amount of HDL-cholesterol [41]. The role of *RUNX1* in promoting HDL levels in mice is consistent with our finding that higher expression of *RUNX1* is associated with higher HDL levels in humans.

Module significance analysis links *RUNX1* to cholesterol production via interactions with *SREBF2*. A physical PPI module containing *RUNX1* and *SREBF2* is significantly enriched for genes with low p-values for association with HDL (Fig 3b). *RUNX1* is connected to *SREBF2* via *CEBPB* and *SRF* (Fig 3b). *SREBF2* upregulates genes involved in cholesterol production, including HMG-CoA reductase, the target of the cholesterol-lowering statin drug family. Statins suppress lower cholesterol by inhibiting HMG-CoA reductase [42, 43]. Interestingly, an intronic variant (rs2834707) in *RUNX1* has been previously associated with HDL in the GWAS Catalog [37]. However, there is no prior association of *RUNX1* expression with HDL in TWAS Atlas [36].

**Fig 3:**
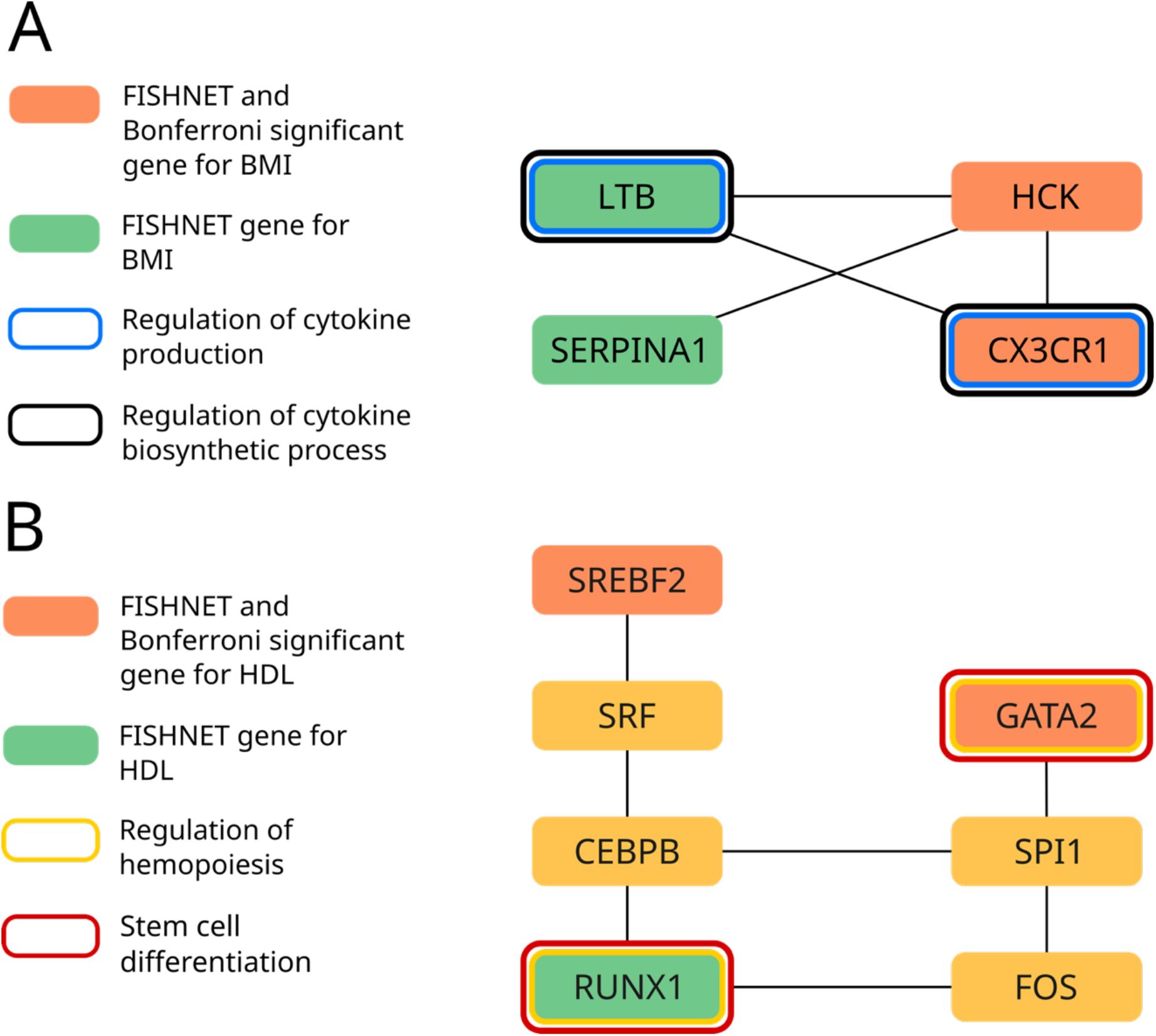
FISHNET genes interact with genome-wide significant genes in significant network modules. (A) A significant co-expression module for BMI contains 4 replicated FISHNET genes. Two of these (*HCK* and *CX3CR1*) are Bonferroni-significant in LLFS and two are not (*LTB* and *SERPINA1*). *LTB*, which is not genome-wide significant, directly interacts with *CX3CR1* and *HCK* and, like *CX3CR1*, participates in cytokine production and cytokine biosynthetic processes. (B) A significant InWeb PPI module for HDL contains a replicated FISHNET gene (*RUNX1*) with 2 Bonferroni-significant and replicated FISHNET genes (*GATA2, SREBF2*). *RUNX1* participates with *GATA2* in the regulation of hemopoiesis and stem cell differentiation. *RUNX1* is connected to both *SREBF2* and *GATA2* via two mediator genes.

### FISHNET genes replicate at a higher rate across p-value and FDR thresholds

We compared the replication rate of all genes satisfying different p-value thresholds with the replication rate of FISHNET genes above the same thresholds. In the summary statistics from TWAS of 11 cardiovascular risk traits in the LLFS cohort, the FISHNET genes had a higher replication rate than all genes satisfying the same p-value thresholds (Fig. 4a). Specifically, we set a series of p-value thresholds, starting with the genome-wide, Bonferroni-corrected significance threshold (p ≤ 3.5 × 10^−6^) and increasing by factors of 10 until we reached p ≤ 3.5 × 10^−2^. Genes were categorized into cumulative bins such that each bin includes all genes satisfying its threshold as well as those satisfying any lower thresholds. The replication rate of FISHNET genes at p ≤ 3.5 × 10^−5^ (53.4%), a factor of ten more liberal than the Bonferroni criterion, was similar to that of all Bonferroni significant genes (56.6%). More generally, the replication rate of FISHNET genes at each threshold was similar to that of non-FISHNET genes at a one-order-of-magnitude more stringent threshold, a trend observed across all thresholds up to p ≤ 3.5 × 10−^2^ (Fig 4a). The same analysis in 23 GWAS discovery datasets from the 2019 Dream Challenge showed the same trend (Fig 4b). Among genome-wide significant genes (p <= 3.5 × 10^−6^), those that were also FISHNET genes had a higher replication rate than the rest. We also compared the replication rate of all genes satisfying different FDR thresholds with that of FISHNET genes satisfying the same thresholds (Fig 4c, Fig 4d). In the LLFS data, FISHNET genes replicated at a rate similar to all genes satisfying smaller by a factor of four (2 bars in Figure 4c). The result from GWAS gene-level summary statistics are even stronger (Figure 4d). The evidence from both datasets suggests relaxing the FDR threshold by a factor of 4 for FISHNET genes will not compromise the replication rate.

**Fig 4:**
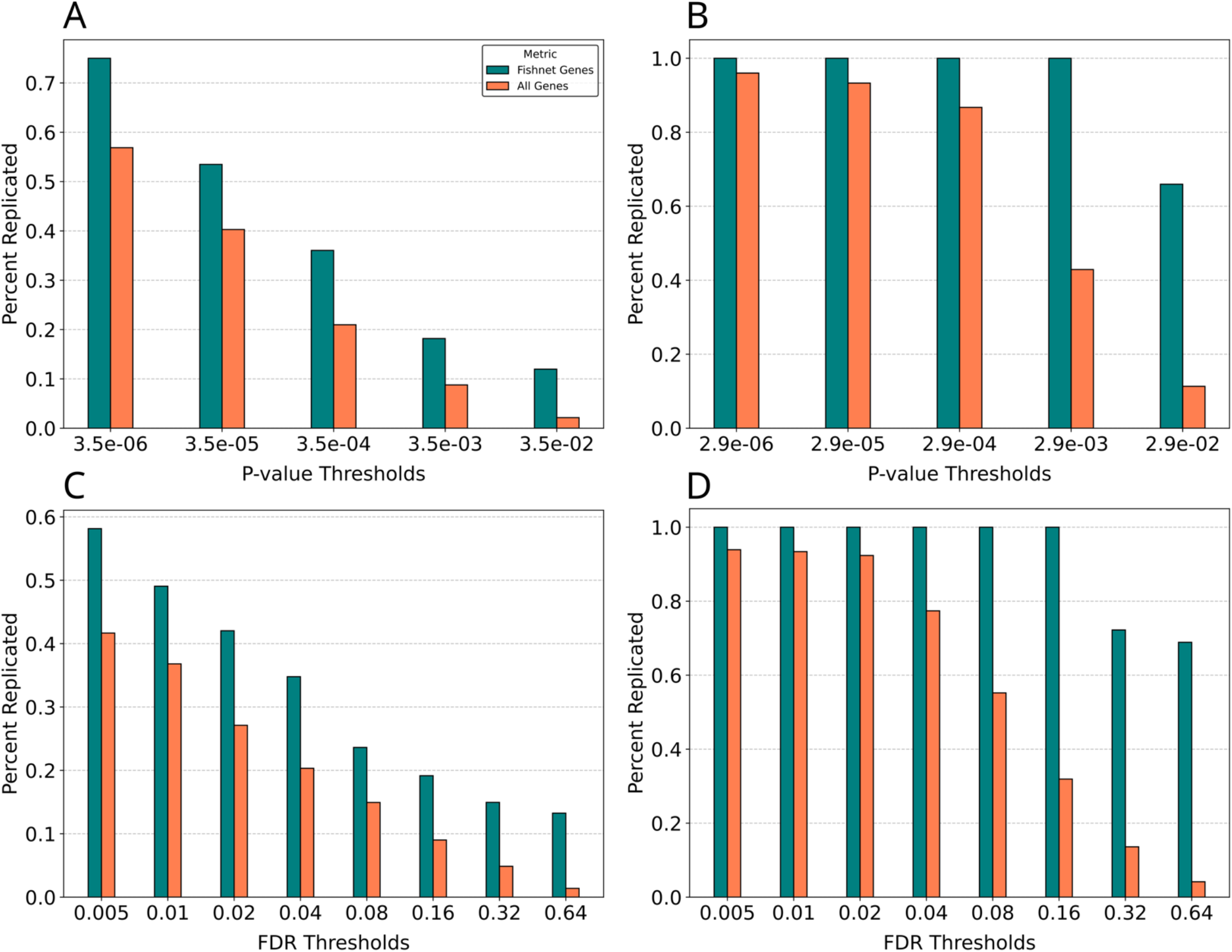
Replication rate across P-value and FDR thresholds. The X-axis shows different p-values and FDR thresholds. The y-axis shows the percentage of replicated genes within the corresponding replication set at a given threshold. (A) shows the replication rate across p-value thresholds in the LLFS cohort (genome-wide significant threshold: p ≤ 3.5 × 10^-6^). (B) shows the replication rate across p-value thresholds in the GWAS summary datasets (genome-wide significant threshold: p ≤ 2.9 × 10^-6^). (C) and (D) show the replication rate across FDR thresholds in the LLFS cohort and GWAS summary datasets, respectively. Across all p-values and FDR thresholds, FISHNET has a better replication rate compared to all genes satisfying the respective thresholds.

### FISHNET is not suitable for co-expression networks built from the same expression data used to generate the p-values

The intuition behind FISHNET is that, under the null hypothesis, gene p-values should be randomly distributed across the network. We hypothesized that this might not be the case when the gene p-values come from association of gene expression levels with traits and the network is generated from the same gene expression data. The reason is that genes in modules of co-expression networks are expected to have similar expression patterns in the data used to generate the network. If the same data are used for association with traits, genes with similar expression patterns across participants can be expected to have similar p-values. Therefore, genes in the same module may have similar p-values, even when those p-values are large and the genes are therefore unlikely to be associated with the trait. To test this hypothesis, we built a co-expression network from the 1810 LLFS gene expression samples by using GENIE3 (https://github.com/vahuynh/GENIE3) [44] (LLFS-net). Modules were identified using the K1 method and submitted to FISHNET, along with p-values for association of the same gene expression levels with 11 cardiovascular risk traits. Across the 11 traits, FISHNET identified 162 enriched module-trait pairs, compared to 38 for the co-expression network based on independent GEO data. A histogram of p-values across modules showed a U-shaped distribution with substantial over-representation of p-values near 1.0 (Fig. 5a), consistent with high P-values clustering in network modules. The same histogram from the GEO network showed much less over-representation or large p-values (Fig 5b). To further investigate this, we reversed the ranking of genes by p-values, so that the most significant genes had the largest p-values and the least significant ones had the smallest p-values. Genes that are not truly associated with the trait, which have now been reassigned the smallest p-values, should not cluster into modules and so no significant modules should be found.

**Figure 5:**
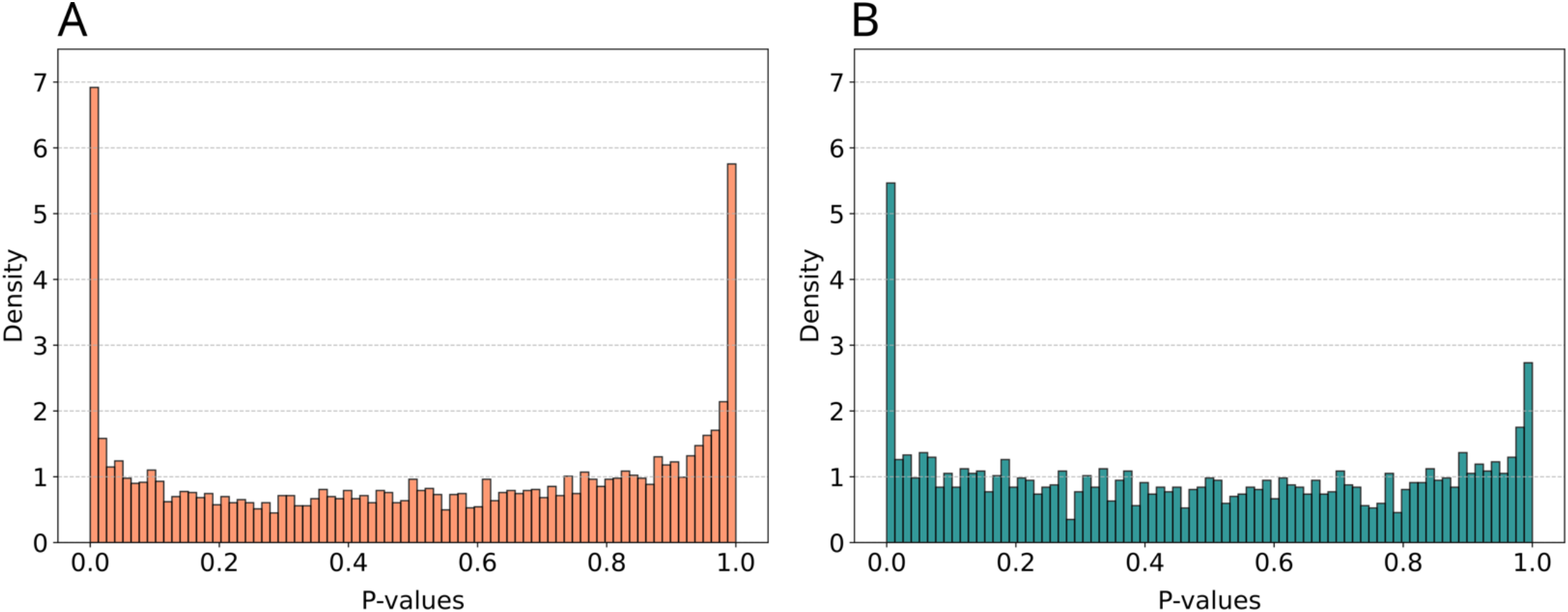
Comparison of module p-value distributions across two types of co-expression networks using LLFS TWAS summary statistics as input. (A) shows the distribution of module p-values from the LLFS co-expression network. The distribution has a prominent U-shape in this case. (B) shows the distribution of module p-values from the GEO co-expression network.

However, when the modules were from LLFS-net, 5 significant modules were found after p-value reversal; when they were from the GEO network built from independent data, no significant modules were found. FISHNET software outputs the number of significant modules after rank reversal as a diagnostic evaluation of the modules’ suitability for the summary statistics.

### An alternative thresholding mechanism allows more control over FISHNET FDR

FISHNET only outputs a gene set if it can identify one that meets its internal thresholds on permutation-based FDR and quantile (Fig. 1). To search for such a set, it first identifies the set *A*_0_ of all genes that (1) are in a module that is significantly enriched for low p-values and (2) are annotated with a biological process term enriched among genes in that module. It then intersects *A*_0_ with *B*_*k*_, the set of all genes with the k lowest p-values, where the default value of k is 5% of the total number of genes, and tests *A*_0_⋂*B*_*k*_ to see whether it satisfies the FDR and quantile criteria. In all but one of the FISHNET runs reported above in which *A*_0_ is not empty, *A*_0_⋂*B*_*k*_ does satisfy the criteria. But when it does not, FISHNET then tests *A*_0_⋂*B*_*k*−10_, *A*_0_⋂*B*_*k*−20_, … until if finds an intersection that satisfies the criteria. In one of the runs described above, this process was necessary and resulted in a satisfactory intersection, *A*_0_⋂*B*_70_. Figure 6A shows the empirical FDR as a function of the percentage of genes included in *B* for this trait, male waist circumference, and in this case the line is monotonically decreasing as the *B* set gets smaller and smaller. However, there is no guarantee that it will – Figure 6A also shows the FDR line for triglycerides, which bounces up and down with no significant trend. It is therefore possible that the FDR could fail for all values of the *B* set. In order to increase the likelihood of identifying a non-empty subset of *A*_0_ that satisfies the criteria, we implemented an alternative approach.

**Figure 6:**
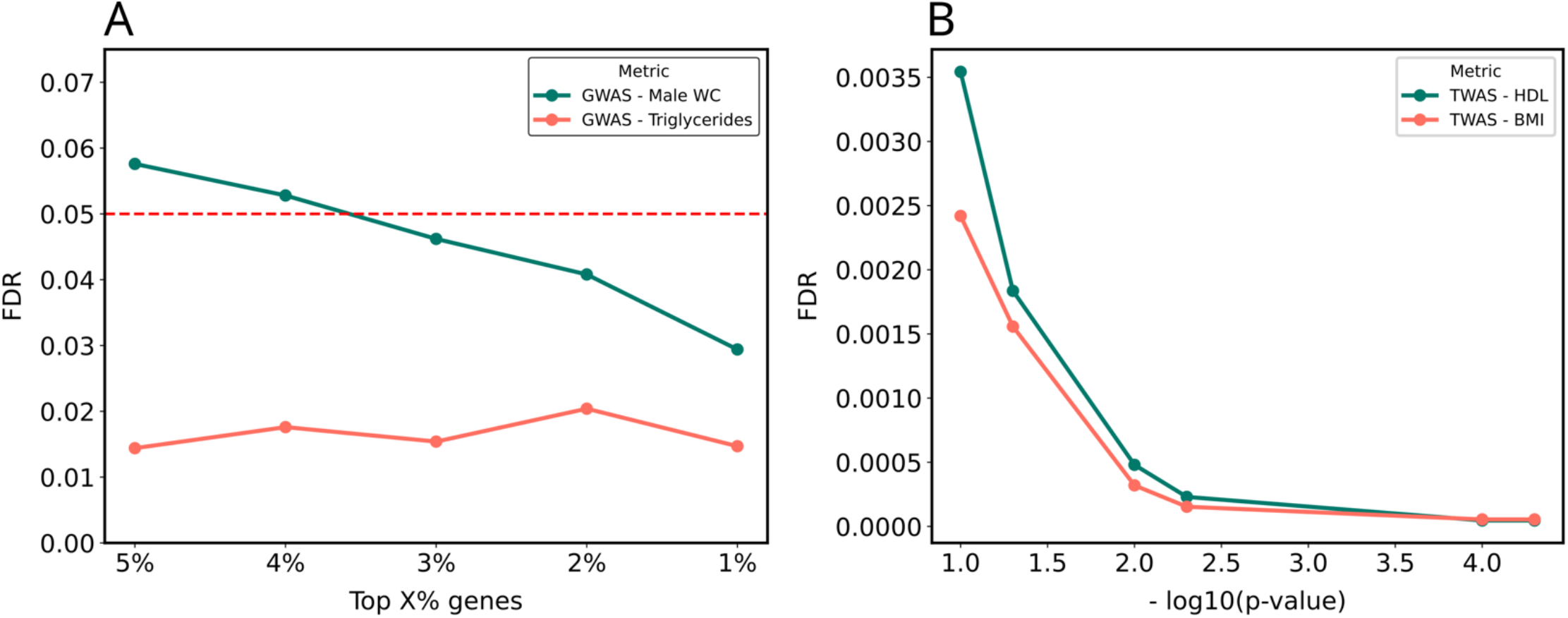
(A) The change in FDR as a function of top X% genes selected in the thresholding mechanism based on iteratively removing genes with the highest p-values for GWAS on male waist circumference and triglycerides levels. (B) shows the change in FDR as a function of module p-values in the thresholding mechanism based on iteratively removing modules based on module significance p-values (module-based filter) for TWAS on BMI and HDL.

In the alternative approach, instead of removing genes with the highest p-values, we iteratively excluded modules based on module significance p-values (module-based filter, Supplementary Figure 1). Starting with modules that had Bonferroni-adjusted p-values ≤ 0.1, the candidate FISHNET gene set was iteratively reduced to genes within modules meeting increasingly stringent thresholds until the FDR and percentile criteria were met. Notably, using the module-based filter decreased FDR monotonically as module p-value thresholds became more stringent (Fig 6b). On the LLFS TWAS summary statistics, this alternative method performed comparably to the original FISHNET pipeline in terms of replication rates (Supplementary Figure 2a, 2b). On the 23 GWAS summary datasets from Choobdar et al. 2019 [14]. FISHNET genes obtained by the alternative method (module-based filter) showed slightly lower replication rates compared to other genes with the lowest gene p-values while showing substantially higher replication rates for genes with higher p-values.

Overall, iteratively removing genes with high p-values outperformed the module-based filter across two datasets and ensured that FISHNET genes met the FDR and percentile rank criteria, establishing it as the preferred method in this work.

## Discussion

Multiple testing correction approaches such as Bonferroni adjustment [7, 8] and FDR control [9, 10] reduce false positives at the cost of increasing false negatives. FISHNET offers a way to relax significance thresholding while maintaining the replication rate.

Across 11 traits associated with cardiovascular risks in the LLFS cohort, FISHNET identified 34 replicable gene-trait relationships, 19 of which were not genome-wide significant in TWAS according to standard thresholds.

### A robust, multi-purpose tool for gene prioritization

An easy-to-install, easy-to-use implementation of FISHNET is available at https://doi.org/10.5281/zenodo.14765850. Given gene-level summary statistics and modules derived from a gene-gene interaction network, it generates (1) a prioritized list of FISHNET genes, (2) a diagnostic evaluation of the modules’ suitability for the summary statistics, and (3) a list of significant network modules along with enriched GO terms associated with module genes. FISHNET also outputs FDR and quantile of the number of candidate FISHNET genes identified, based on permutation analysis. Users can customize the pipeline by defining the initial threshold for the gene set to be considered for the FISHNET pipeline (default: top 5% genes ranked by p-values). Users can also select between module-based (Supplementary Figure 1) and gene-based (Figure 1) filters.

While we emphasized FISHNET’s ability to identify gene-trait relationships that may be overlooked by traditional association analyses, its value as a tool for identifying significant modules and enriched GO terms does not depend on the choice of gene significance threshold. Users who choose not to relax their significance thresholds can still input gene-level summary statistics to uncover significant modules and their associated enriched GO terms. The broad range of accepted inputs, detailed outputs, and customizable features make FISHNET a robust and versatile tool for network-based gene prioritization and functional analysis.

FISHNET also improves on traditional network-based methods for association analysis in both methodology and evaluation. One of the most popular approaches is to select previously known trait-associated genes as seed nodes and propagate their scores across local neighborhoods in the network to predict new gene-trait relationships [45, 46]. This approach creates a bias against detecting genes that affect the trait via pathways that are different from those of known genes. FISHNET uses all gene-level p-values, which inherently reduces pathway bias, and it identifies modules significantly over-represented for genes with low p-values even if those genes are far from significant when considered individually. Furthermore, FISHNET incorporates GO biological process annotations to identify genes that (1) interact in significant network modules and (2) participate in a common biological process. In terms of evaluation, the most common evaluation metrics to validate network-based approaches are leave-one-out methodology [46, 47] and validation with genes from published drug target databases [48]. These metrics test the methods’ ability to perform well in recovering *known* gene-trait relationships. However, the ultimate goal is to uncover *novel* gene-trait relationships. To achieve this, we evaluated FISHNET by comparing the replication rate of FISHNET genes against those meeting standard Bonferroni or FDR thresholds.

### Findings from FISHNET and recommendations

Our results suggest best practices for using FISHNET. Among the modularization algorithms tested, we recommend the kernel clustering method K1 [14] (see Methods) when prioritizing both reproducibility and replicability. The modularity optimization method, [M2] performed best in replicability, while the random walk approach performed best in reproducibility (Fig 2b). An alternative approach would be to use multiple modularization algorithms and take the union of the FISHNET genes they identify. While we did not evaluate the replicability and reproducibility of FISHNET genes as the function of the number of the modularization algorithms used, all three algorithms tested performed well in at least one evaluation metric. This suggests that combining results from multiple algorithms is a promising strategy.

Among networks, FISHNET performed well on the InWeb physical PPI and GEO co-expression networks but poorly on the STRING functional network (Fig 2a). We recommend using networks based on physical PPI and co-expression interactions over those based on other functional interactions. We also caution against using the same dataset to generate summary statistics and construct gene-based networks, as this can lead to unreliable FISHNET gene-trait relationships (Fig 5a, 5b). Based on our findings (Fig 4a, 4b, 4c, 4d), we recommend using a slightly relaxed Bonferroni-or FDR-based threshold and prioritizing FISHNET genes that meet these relaxed criteria alongside the genes that are significant under the original threshold. This recovers gene-trait relationships missed by standard thresholds while maintaining the replication rate.

### Limitations and opportunities for broader applications

There are many potential use cases for FISHNET that have not yet been validated. First, the gene-level p-values can come from any source, including RNA-Seq experiments (such as comparing cells treated with a drug to untreated cells) and CRISPR screens to identify genes that affect cellular traits [49, 50]. Second, FISHNET has been tested only on specific gene-gene networks, but others might give better, worse, or complementary results. For example, FISHNET was tested on co-expression networks, but these are neutral as to the molecular mechanism that causes each pair of linked genes to have similar expression patterns. A mechanistic alternative is networks that link transcription factors to their direct targets [51-54]. Modules from mechanistic gene regulatory networks could elucidate the specific transcription factors mediating gene–trait relationships. Third, FISHNET has only been tested on modules identified by three algorithms. Since different traits and networks performed optimally with modules from different algorithms, it might be valuable to test a broader range of modularization strategies. More generally, FISHNET uses only the set of genes in each module, not its internal connectivity. Thus, biologically coherent gene sets from any source could be used. For example, the sets of genes directly regulated by each transcription factor (TF) could be used, in which case no modularization algorithm is required. Fourth, the final stage of FISHNET currently uses Gene Ontology Biological Process terms to identify biological functions enriched among genes in significant modules, but gene sets from other sources could be used, too. For example, gene sets defined by drug target discovery databases such as DrugBank [55, 56], DisGeNET [57], and Open Targets [58] [59] could provide complementary insights and reveal new genes. The versatility of our implementation allows users to try out any of these sources for gene-level p-values, module gene sets, and functional gene sets. Future work validating FISHNET with new knowledge sources will greatly expand its applicability.

## Acknowledgments

We are grateful to the entire Long Life consortium, its participants, and its investigators, without whom this work would not have been possible. This work was supported by grant AG063893 from the National Institute on Aging.

The Framingham Heart Study is conducted and supported by the National Heart, Lung, and Blood Institute (NHLBI) in collaboration with Boston University (Contract No. N01-HC-25195, HHSN268201500001I and 75N92019D00031). This manuscript was not prepared in collaboration with investigators of the Framingham Heart Study and does not necessarily reflect the opinions or views of the Framingham Heart Study, Boston University, or NHLBI.

## Data and code availability

The FISHNET pipeline is available at https://doi.org/10.5281/zenodo.14765850. All the summary statistics from association analyses used as inputs, and the FISHNET outputs are available in Supplementary Files. The input datasets to both LLFS and FHS association analyses have not been deposited in public repositories due to data use constraints.

