## Supplementary material for "FISHNET: A Network-based Tool for Analyzing Gene-level P-values to Identify Significant Genes Missed by Standard Methods": Supplementary_Figures.docx

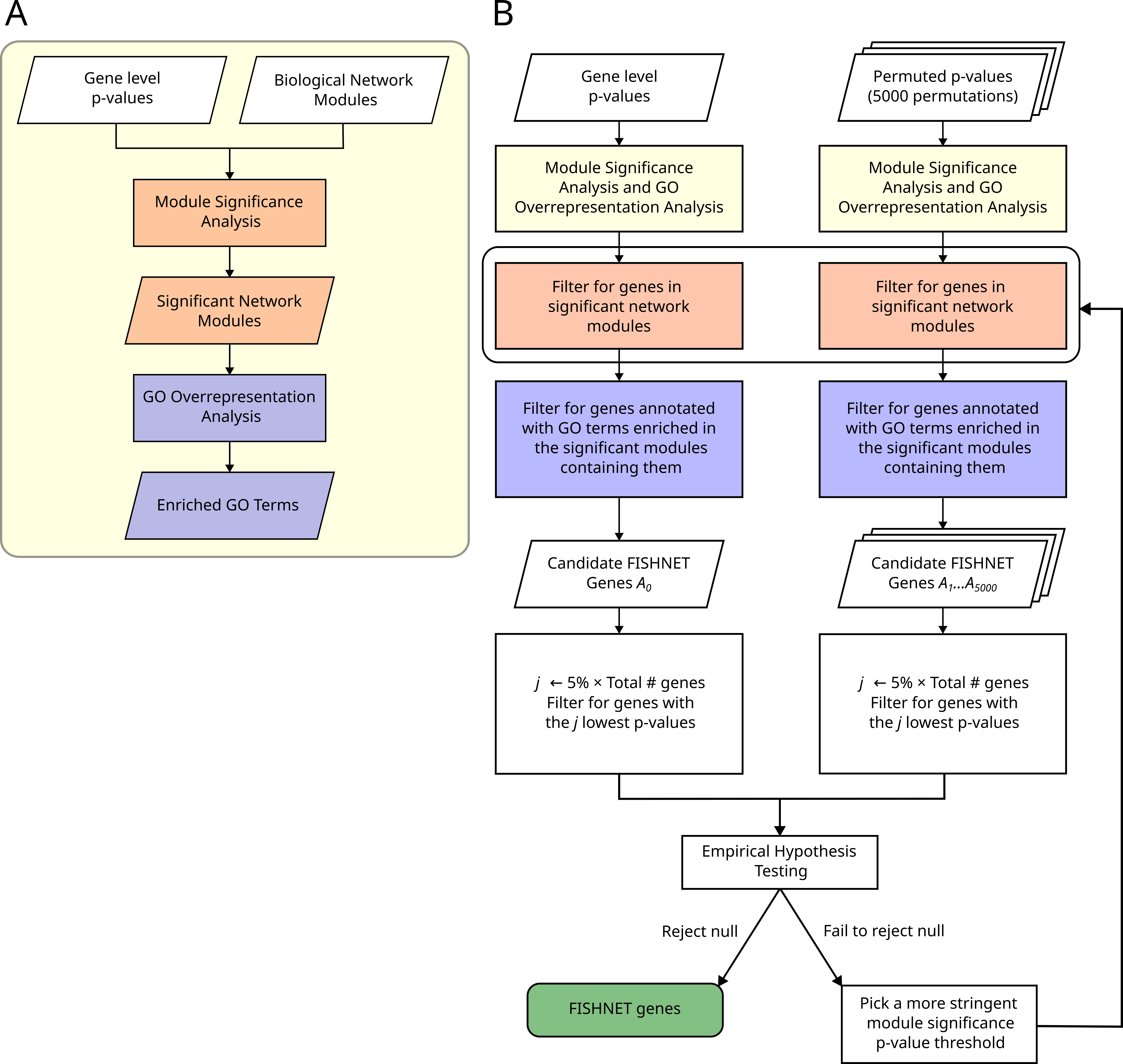
Supplementary Fig 1: The gene-level omics-WAS summary is fed into module enrichment analysis, which is performed by PASCAL. PASCAL outputs significant modules and their p-values. Gene ontology over-representation analysis identifies biological processes with significant over-representation among genes in each significant module. (B) The workflow illustrates the gene prioritization mechanism for identifying FISHNET genes by applying the thresholding based on module p-values. When the null hypothesis is rejected, a more stringent module p-value is picked iteratively, and filters are re-applied to identify candidate FISHNET genes for empirical hypothesis testing.

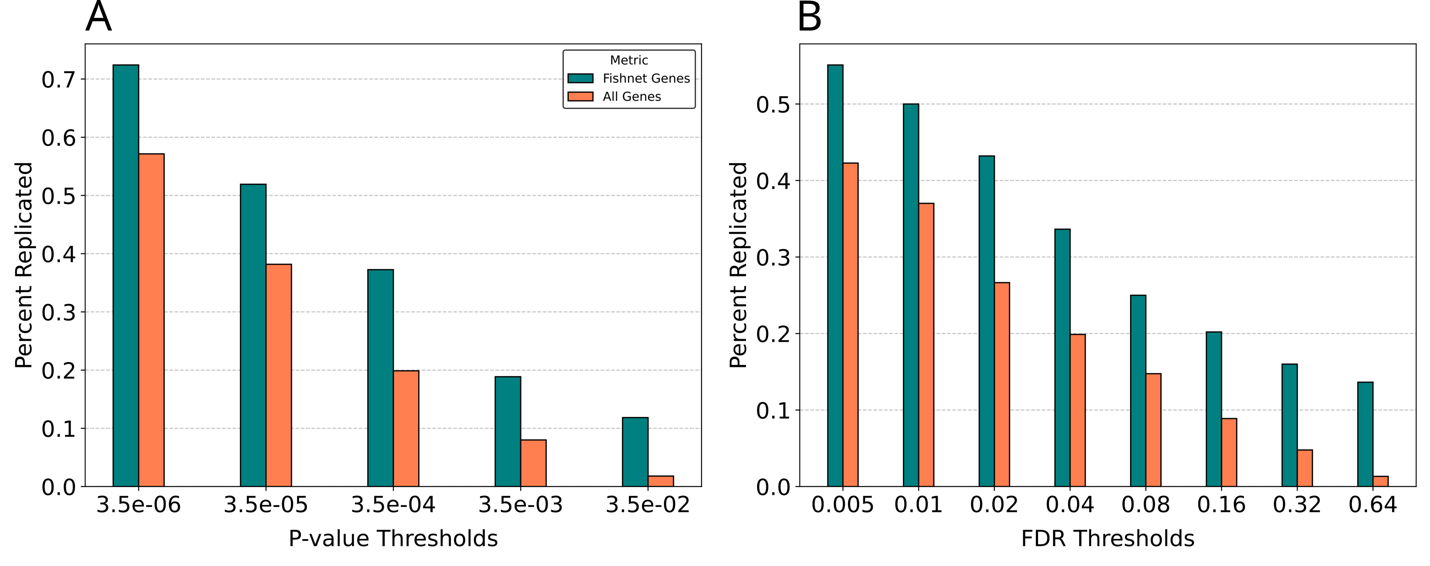

Supplementary Fig 2: Replication rate across P-value and FDR thresholds using the thresholding based on module p-values. The X-axis shows different p-values and FDR thresholds. The y-axis shows the percentage of replicated genes within the corresponding replication set at a given threshold. (A) shows the replication rate across p-value thresholds in the LLFS cohort (genome-wide significant threshold: p ≤ 3.5 × 10^-6^). (B) shows the replication rate across FDR thresholds in the LLFS cohort).

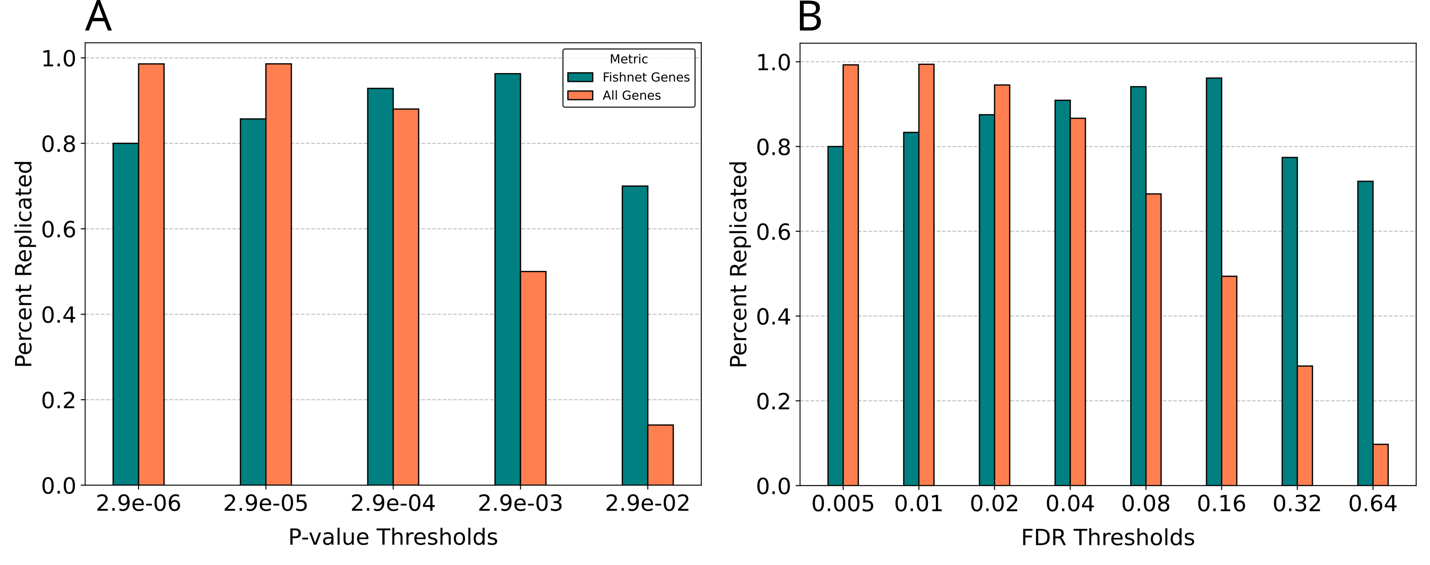

Supplementary Fig 3: Replication rate across P-value and FDR thresholds using the thresholding based on module p-values. The X-axis shows different p-values and FDR thresholds. The y-axis shows the percentage of replicated genes within the corresponding replication set at a given threshold. (A) shows the replication rate across p-value thresholds in the GWAS summary datasets (genome-wide significant threshold: p ≤ 2.9 × 10^-6^). (B) shows the replication rate across FDR thresholds in the GWAS summary datasets.
